# UHRF1-SRA recognizes symmetric non-CG methylated DNA through dual-flip out of 5-methyl cytosines

**DOI:** 10.1101/2019.12.17.880419

**Authors:** Naveen Kumar Nakarakanti, Suman Abhishek, Waghela Deeksha, Eerappa Rajakumara

**Affiliations:** Macromolecular Structural Biology Lab, Department of Biotechnology, Indian Institute of Technology Hyderabad, Kandi, Sangareddy, Telangana-502285, India

**Author notes:** Corresponding author Eerappa Rajakumara, Associate Professor, Macromolecular Structural Biology Lab, Department of Biotechnology. **Equal contribution:** These authors have contributed equally to the manuscript (Naveen Kumar Nakarakanti, Suman Abhishek).

## Abstract

Non-CG DNA methylation (non-mCG) is enriched in the genome of brain neurons and germline cells. Non-mCG is differentially distributed on neuronal X-chromosome in males and females. Accumulation of non-mCG during postnatal brain development correlates with reduced gene expression and inactivation of distal regulatory elements, and allele specific gene regulation. Recently, UHRF1 has been found to contribute to *de novo* non-CG methylation, however, whether UHRF1 could recognize non-mCG is not known. Here, we have demonstrated through calorimetric measurements that the SRA domain of UHRF1 can recognize mCH and fully-mCHG, types of non-mCG. Furthermore, our ITC binding analyses with methylated CG DNA (mCG) revealed 6-fold decrease in binding affinity for fully-mCG compared to hemi-mCG and, despite symmetrical 5mCs, stoichiometry of 1:1 for UHRF1 SRA binding to fully-mCG indicates UHRF1 may not form stable complex with fully-mCG DNA. In contrast, UHRF1 SRA recognizes fully-mCHG with a stoichiometry of 2:1 protein to DNA duplex, and has tighter binding compared to fully-mCG. Crystal structure of UHRF1 SRA bound to 5mC containing DNA in fully-mCHG context revealed dual flip-out mechanism of 5mC recognition. Altogether, this study indicates that UHRF1 SRA also recognizes non-mCG DNA, besides known hemi-mCG DNA and exhibits contrasting mechanisms for hemi-mCG and fully-mCHG DNA recognition. These findings may open a new window to investigate the biological function of non-CG methylation recognition by the UHRF1.

## Introduction

The methylation of cytosine (5-methyl cytosine: abbreviated as 5mC or mC) in the DNA at CG (mCG) sites in the mammalian genome is an epigenetic modification, locally and globally regulating gene expression, that is involved in various cellular processes, such as cell differentiation, transposon silencing, genomic imprinting, and carcinogenesis. DNA methylation in non-CG contexts (non-mCG), either asymmetric or symmetric, is present in almost all human tissues and correlated with tissue-specific functions(Schultz et al. 2015). mCH (where H is A, T or C) is an asymmetric non-mCG, enriched in two main categories of the mammalian cells: brain neurons (Lister et al. 2013; Xie et al. 2012)and germline cells. Embryonic stem cells (ESCs) (Lister et al. 2009), induced pluripotent stem cells (iPSCs)(Lister et al. 2011), oocytes(Guo et al. 2014; Okae et al. 2014; Shirane et al. 2013)and primordial germ cells(PGCs)(Gkountela et al. 2015; Guo et al. 2015; Kobayashi et al. 2013; Seisenberger et al. 2012; Tang et al. 2015) are mCH-enriched germline cells. Presence of mCH in myocytes is a proof for non-CG methylation in somatic cells(Schultz et al. 2015). mCH is a distinct feature of neuronal epigenome that is differentially distributed on X chromosome in males and females(Keown et al. 2017; Mo et al. 2015). In addition, millions of CA and CT non-CG methylations accumulate during postnatal brain development in mice(Lister et al. 2013; Xie et al. 2012)and it correlates with reduced gene expression and inactivation of distal regulatory elements in a highly cell type-specific manner (Mo et al. 2015). A recent study has shown that functioning of mCH in neuron is independent of mCG, and dynamic nature of mCH is correlated with allele specific gene regulation(Keown et al. 2017). Fully-mCHG (5′-mCHG-3′/3′-GNmC-5′: where ‘H’ is A, T or C and ‘N’ is T, A or G), asymmetric non-CG methylation, is highly prevalent in plants. There are studies reporting the presence of symmetric mCWG (W is A or T), a class of fully-mCHG, on the promoters of specific genes in mammalian cell lines(Malone et al. 2001). Comparison of mCH across different human cell types has demonstrated the presence of symmetric mCHG in TACAG sequence context in ESCs(Chen et al. 2011).

DNA methylation patterns are transmitted with high fidelity during DNA replication. In mammals, CG and CH DNA methylation is established by the DNA methyltransferase 3 (DNMT3) family of *de novo*methyltransferases and CG DNA methylation is maintained by the DNMT1 (Cheng and Blumenthal 2008; Goll and Bestor 2005; Kim et al. 2009; Ramsahoye et al. 2000). DNMT1 shows a strong preference for CG hemi-methylated DNA (hemi-mCG: 5′-mCG-3′/3′-GC-5′), and it is recruited to the DNA replication fork through direct interactions with PCNA (proliferating cell nuclear antigen)(Bostick et al. 2007; Chuang et al. 1997; Hermann et al. 2004). Proteins that contain SET- and RING-associated (SRA) domains have been demonstrated to play unequivocal role at the level of establishment and/or maintenance of both CG and CH DNA methylation in plants and mammals(Bostick et al. 2007; Johnson et al. 2008; Kraft et al. 2008; Sharif et al. 2007; Woo et al. 2008). UHRF1 (ubiquitin-like, containing PHD and RING finger domains 1, also known as Np95 and ICBP90)recognizes hemi-mCG sites via an SRA domain through a flip-out of 5mC from the duplex DNA, and recruits DNMT1 to these sites(Arita et al. 2008; Avvakumov et al. 2008; Bostick et al. 2007; Hashimoto et al. 2008). One of the two loops of UHRF1 SRA (Fig. 1A), which hand grasps the hemi-mCG DNA duplex, is predicted to clash sterically if a symmetric methylated cytosine is present on the complementary strand of the DNA. Consistent with the structure of the UHRF1 SRA bound to hemi-mCG duplex DNA, binding affinity of UHRF1 SRA to hemi-mCGis stronger compared to fully-mCG(Bostick et al. 2007).UHRF1 has also been reported to participate in inter-stand cross links repair (ICLs), through specifically interacting with ICLs *in vivo* and *in vitro* and recruits FANCD2 to initiate Fanconi Anemia DNA repair pathway (Liang et al. 2015; Tian et al. 2015).

**Fig. 1:**
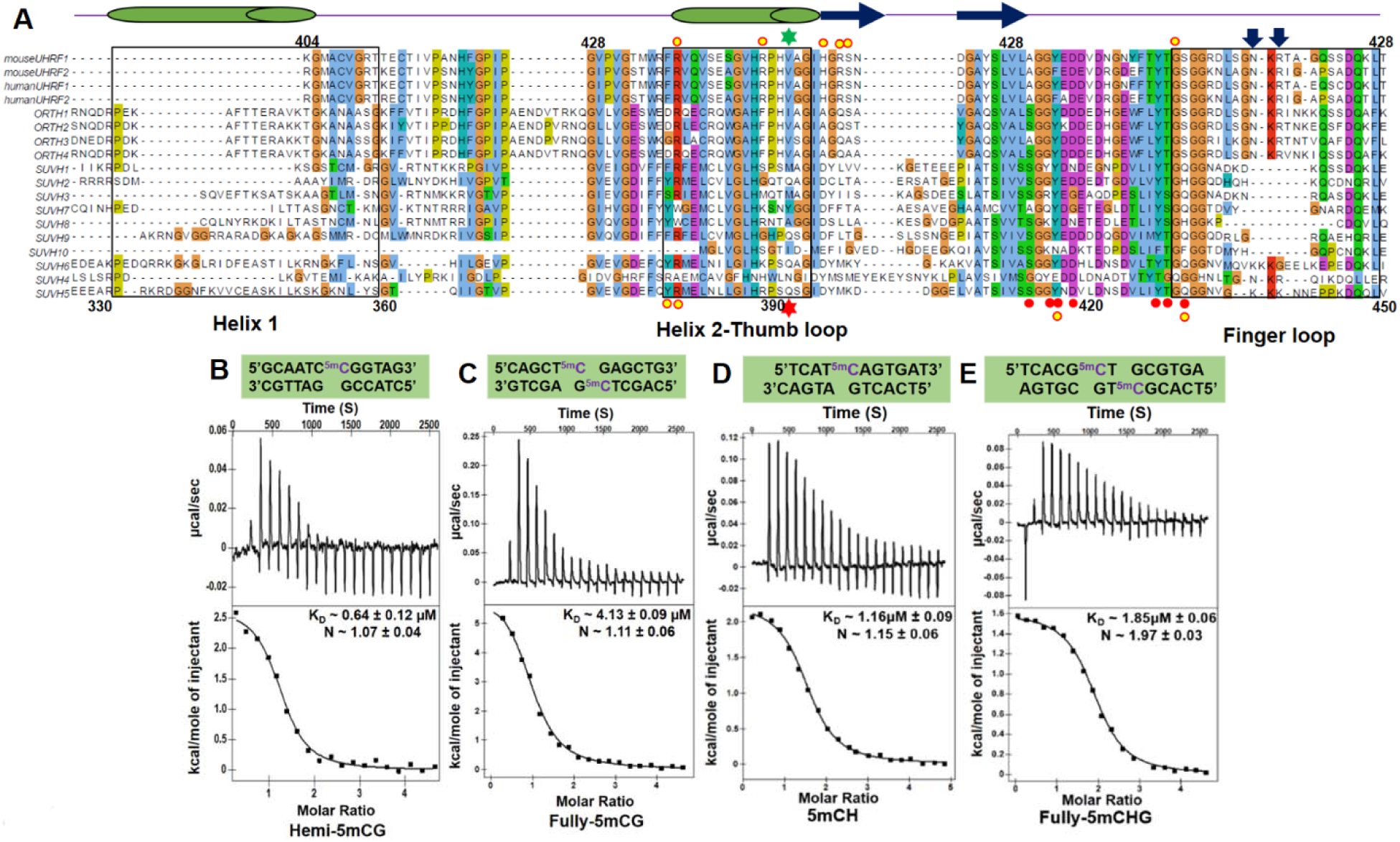
Multiple sequence alignment of SRA domains of different proteins and ITC studies of UHRF1 SRA binding to 5mC-containing DNA in different sequence contexts. (A) Multiple sequence alignment: UHRF proteins numbering is based on mouse UHRF1 at the top and SUVH family is based on SUVH5 SRA at the bottom. The secondary structural elements are indicated above the sequence (α-helices are in green cylinders, β-strands are in blue arrows, and disordered regions are represented by simple blue line). The helix-1, helix-2, thumb and finger loop regions are highlighted with a rectangle. Residues are highlighted with default Clustal X colouring. Inverted arrows in blue correspond to residues that interact with the orphaned guanine and adjacent cytosine pair and masking the unmodified cytosine in the UHRF1 SRA. Green and red stars represents the residues of UHRF1 SRA and SUVH5 SRA, respectively, that inserts into the DNA duplex. Red circles designate residues of the SRA domain of UHRF1 and SUVH5 that interact with 5mC in their respective binding pockets. Yellow circles designate residues ofUHRF1 SRA and SUVH5 SRA that interact with the DNA backbone. ITC measurements of the binding of the UHRF1 SRA domain to **(B)** hemi-mCG DNA **(C)** fully-mCG DNA **(D)** mCH DNA **(E)** fully-mCHG DNA. The DNA sequence used for ITC studies are mentioned above thermograms. Experimental details are provided in the Materials and Methods.

However, there is no data available reporting recognition of non-mCG DNA by the UHRF1 SRA. Our binding study establishes that UHRF1 SRA recognizes fully-mCG (5′-mCG-3′/3′-GmC-5′) with a stoichiometry of ∼1, which indicate, despite two symmetrical 5mCs, one UHRF1 SRA recognizes one fully-mCG duplex DNA. Our studies are the first to establish that UHRF1 SRA recognizes mCH and fully-mCHGDNA. The recognition of non-CG methylated DNA is supported by the structure of UHRF1 SRA bound to fully-mCHG duplex DNA. Structure of UHRF1 SRA bound to fully-mCHG DNA reveals two molecules of UHRF1 SRA recognize a fully-mCHG duplex DNA through dual flip-out of symmetric 5mCs that is consistent with stoichiometry of ∼2 (2 UHRF1 SRA: 1 fully-mCHG) estimated by the ITC binding studies. Importantly, this study unearths the non-CG methylated DNA recognition by the UHRF1 SRA, which supports the involvement of UHRF1 in *de novo* non-CG methylation (Maenohara et al. 2017).

## Results

### Calorimetric studies of CG methylated DNA binding specificity of UHRF1 SRA domain

UHRF1 SRA binds both hemi- and fully-mCG DNA, and it exhibits higher preference for hemi-mCG (Arita et al. 2008; Avvakumov et al. 2008; Bostick et al. 2007; Hashimoto et al. 2008) (Fig. 1B-C), however no data is available on binding stoichiometry. Here, we used isothermal titration calorimetric (ITC) approach to investigate binding affinity and stoichiometry for UHRF1 SRA and DNA duplexes in fully- and hemi-methylated CG contexts. The UHRF1 SRA domain binds to hemi-mCG with a molar dissociation constant (K_D_) of 0.64 µM (Fig. 1B) with stoichiometry of ∼1. Consistent with the previously reported EMSA study, the binding affinity decreased by a factor of 6.4 (K_D_ = 4.13 µM) (Fig. 1C) for fully-mCG compared to mCG (Bostick et al. 2007). Recognition of mCG DNAs in both methylation statuses is entropy driven with ΔS of 36.98 and 46.25 cal.mol^-1^K^-1^ for hemi- and fully-mCG duplexes, respectively (Table 1). Surprisingly, UHRF1 SRA domain exhibits stoichiometry of ∼1 for fully-mCG DNAs, which reflects an SRA domain bound per fully-mCG DNA duplex.

**Table 1:** Thermodynamic data and dissociation constant values for the binding of CG and non-CG methylated DNAs by UHRF1 SRA.

| **Modified DNA duplex** | **N (Stoichiometry)** | **$K_D$ (μM)** | **$\Delta H$ (Kcal.mol<sup>-1</sup>)** | **$\Delta S$ (cal.mol<sup>-1</sup>K<sup>-1</sup>)** |
| --- | --- | --- | --- | --- |
| mCG | 1.07 ± 0.04 | 0.64 ± 0.12 | 2.67 ± 0.51 | 36.98 |
| mCH | 1.15 ± 0.06 | 1.16 ± 0.09 | 2.25 ± 0.09 | 33.85 |
| Fully-mCHG | 1.97 ± 0.03 | 1.85 ± 0.06 | 1.61 ± 0.04 | 32.04 |
| Fully-mCG | 1.11 ± 0.06 | 4.13 ± 0.09 | 6.23 ± 0.16 | 46.25 |

### UHRF1 SRA recognizes non-CG methylated DNA

We performed ITC binding studies to investigate whether UHRF1 SRA recognizes non-CG methylated DNAs in the context of mCH and fully-mCHG. Binding affinity was marginally decreased by a factor of 1.8 (K_D_ = 1.16 µM) (Fig. 1D) for mCH and a factor of 2.9 (K_D_ = 1.85 µM) (Fig. 1E) for fully-mCHG compared to hemi-mCG (Fig. 1B). Conversely, UHRF1 SRA shown higher affinity towards mCH and fully-mCHG DNAs compared to fully-mCG DNA. Vis-à-vis comparison reveals that UHRF1 SRA binds to methylated DNA duplexes with increased affinity in the order of fully-mCG, fully-mCHG, mCHand hemi-mCG (Fig. 1). Interestingly, stoichiometry of binding is ∼2 (2 UHRF1 SRA: 1 duplex DNA) for fully-mCHG DNA recognition compared to fully-mCG which has contrasting stoichiometry of ∼1. However, stoichiometry of binding is ∼1 for mCH DNA which is similar to hemi-mCG DNA. Binding of UHRF1 SRA to non-mCGs is also entropically driven (Table 1).

### Structure of the UHRF1 SRA domain bound to fully-mCHG DNA

We have solved the crystal structure of the UHRF1 SRA domain bound to a 13 base pair length duplex DNA containing centrally positioned fully-methylated CHG steps (Fig. 2A). Single crystal of this complex (space group P2_1_2_1_2_1_) was diffracted to 3.0Å resolution (crystallographic statistics listed in Table 2). The structure of the UHRF1 SRA-fully-mCHG complex contains two SRA molecules bound per DNA duplex, with both the symmetric 5mC bases flipped out of the DNA helix and positioned in the binding pockets of individual SRA domains (Fig. 2B). Two SRA molecules bound per duplex is consistent with a stoichiometry of ∼2 observed in ITC binding studies for complex formation of theUHRF1 SRA domain with fully methylated CHG DNA (Fig.1E). Notably, Val451 of thumb-loop from both SRA domains, inserts into the duplex DNA from minor groove, and partly substitute 5mC position in the duplex DNA, however it doesn’t interact with the orphan guanine G5. Thumb-helical region non-specifically interacts with the -1 (G5) that reinforce the complex formation. The flipped out 5mC is positioned in a pocket within the SRA domain such that it is anchored in place via stacking interactions with Tyr471 and Tyr483 and by intermolecular hydrogen bonds between its Watson-Crick edge and the side chain of Asp474, Thr484, the backbonecarbonyl group of Thr484 and the backbone amide nitrogens of Ala468, Gly469 (Fig. 2C).

**Table 2:** X-ray diffraction data and structure refinement statistics. The values for the highest resolution shell is shown in parentheses.

| **Crystal** | **UHRF1 SRA-fully-mCHG** |
| --- | --- |
| Wavelength | 1.54179Å |
| Resolution range | 29.89Å - 3.0Å (3.107Å - 3.0Å) |
| Space group | P 2<sub>1</sub> 2<sub>1</sub> 2<sub>1</sub> |
| Unit cell (Å) | a = 55.9Å, b = 74.8Å, c = 141.5Å |
| $\alpha$ $\beta$ $\gamma$ (°) | $\alpha = \beta = \gamma = 90^\circ$ |
| R<sub>sym</sub> | 0.05297 (0.4682) |
| Completeness (%) | 99.73 (99.92) |
| I/$\sigma$ (I) | 9.78 (1.66) |
| Redundancy | 2.0 (3.5) |
| Number of unique reflections | 12420 (1240) |
| Reflections used for R-free | 621 (61) |
| R work (%) | 0.2459 (0.3728) |
| R free (%) | 0.3040 (0.4350) |
| Number of non-H atoms | 3450 |
| Protein | 2922 |
| DNA | 528 |
| Water | - |
| Protein residues | 390 |
| **R.M.S. Deviation** |  |
| R.M.S.D bond lengths (Å) | 0.006 |
| R.M.S.D bond angles (°) | 1.05 |
| **Mean B values ($\text{\AA}^2$)** | 61.68 |
| Protein | 56.54 |
| DNA | 72.54 |
| Ramachandran favoured (%) | 94.21 |
| Ramachandran allowed (%) | 5.53 |
| Ramachandran outliers (%) | 0.26 |
| Rotamer outliers (%) | 0.00 |

**Fig. 2:**
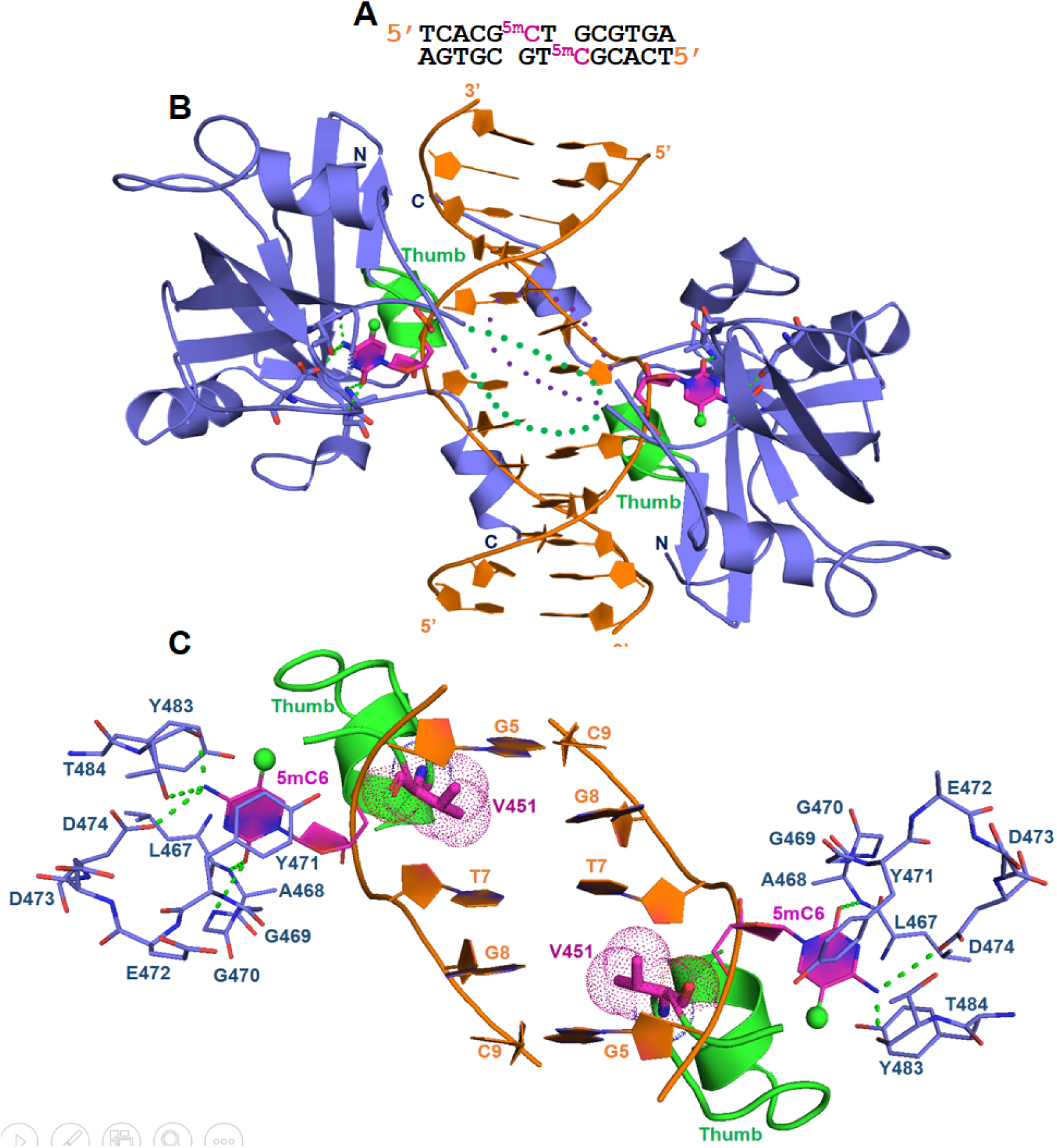
Crystal structure of the UHRF1 SRA domain bound to the fully-mCHG DNA. **(A)** Sequence of the 13-mer self-complementary fully-5mCHG DNA. 5mC base is coloured magenta in both the DNA strands. (**B**) Cartoon (protein and DNA) representation of the 3.0Å crystal structure of UHRF1 SRA complexed with fully-mCHG DNA. The 5mC residues from the complimentary strands are flipped out and are positioned in the binding pockets of the individual SRA domains. The DNA is coloured orange except for 5mC, which is in magenta. The methyl group on cytosine is shown as green sphere. The protein domains are coloured blue. The thumb loop and associated helix is coloured green. (**C**) Magnified view of the base flip out from the duplex. Val451 from the thumb loop is inserted into and fills the gap created by the flipped-out 5mC base. The 5mC base is held between the aromatic rings of Tyr471 and Tyr483 through pi-stacking. Side chain carboxyl group of Asp474, main chain carboxyl of Thr484 and main chain amide group of Ala468 and Gly470 stabilizes the 5mC base in the SRA binding pocket through H-bond interaction.

## Discussion

SRA domain of the plant-specific SUPPRESSOR OF VARIEGATION 3-9 HOMOLOG (SUVH) family H3K9 methyltransferases (MTases) have been reported to recognize CG- and non-CG methylated DNA (Du et al. 2014; Li et al. 2018; Rajakumara et al. 2011). However, SRA domain containing proteins of mammals have been reported to recognize only methylated and hydroxyl-methylated DNA in CG sequence context (Arita et al. 2008; Avvakumov et al. 2008; Bostick et al. 2007; Hashimoto et al. 2008; Zhou et al. 2014). Till date UHRF1 SRA is considered as a bonafide reader of hemi-mCG DNA during replication coupled maintenance of CG methylation in the genome. Our binding and structural studies established that UHRF1 SRA also recognizes non-mCG DNAs, mCH and fully-mCHG. Thus, our studies support the critical role of UHRF1 in maintenance of non-CG methylation in mammalian genome (Maenohara et al. 2017). Binding study analysis revealed binding of one SRA molecule per hemi-mCG and mCH DNA duplexes (Fig. 1 B&D). Therefore, we assume that UHRF1 SRA may recognize mCH DNA through flip-out of only 5mC similar to hemi-mCG recognition.

### Single base spacer is required for recognition of fully-methylated cytosine DNA by UHRF1 through dual flip-out mechanism

ITC measurements established an unexpected binding of one UHRF1 SRA domain per fully-mCG duplex DNA, despite fully-mCG has two symmetrical 5mCs (Fig. 1C). Our efforts to crystallize the UHRF1 SRA-fully-mCG complex to unravel the basis for contrasting stoichiometry and mechanistic insights into 5mCs recognition were unsuccessful. We speculate UHRF1 SRA-fully-mCG complex is not stable as recognition of one of symmetric 5mCs by an SRA domain may negatively affect the recognition of other 5mC by another SRA. To know whether UHRF1 SRA domains have steric clashes if they recognize fully-mCG through dual flip-out mechanisms, SRA from UHRF1 SRA-hemi-mCG complex (PDB ID: 2ZO0) was superimposed on the both protomers of UHRF2 SRA in the UHRF2 SRA-hemi-5hmCG complex structure (PDB ID: 4PW6), in which both 5hmC and unmodified C on complementary strand are flipped out. The modeled new complex structure containing two superposed UHRF1 SRAs bound to hemi-5hmC mimics a dual flip-out of 5mC bases by two protomers of UHRF1 SRA (Supplementary figure 1). Following are possible explanations for decrease in binding affinity compared to hemi-mCG and having unexpected stoichiometry for recognition of fully-5mCG where both symmetrical ‘Cs’ are methylated: (A) Finger loop of UHRF1 SRA may have steric clashes with symmetrical SRA, (B) UHRF1 SRA bound to one of the 5mCs is dislodged during the process of other 5mC recognition by another UHRF1 SRA molecule (Supplementary figure 1). Otherwise, UHRF1 SRA is expected to have a stoichiometry of ∼2 (2 SRA: 1 duplex DNA) for fully-mCG DNA duplex recognition similar to SUVH5 SRA recognition mechanism (Rajakumara et al. 2011; Hashimoto et al. 2008; Arita et al. 2008; Avvakumov et al. 2008).

Surprisingly, ITC binding studies reveal that at least single base spacer between symmetric 5mCs is required for recognition with stoichiometry of two SRA bound to a fully-mCHG DNA duplex (Fig. 1E). UHRF1 SRA may recognize fully-mCHG through three possible ways: both 5mCs flip-out like SUVH5 SRA-fully-mCG complex, one 5mC flip-out and no flip-out of another 5mC, or no flip-out of 5mCs. To understand the mechanism of recognition of fully-mCHG DNA by UHRF1 SRA, we initiated a structure based mechanistic study aimed at solving the crystal structure of the UHRF1 SRA in a complex with fully-mCHG DNA duplex. The crystallographic analysis of this complex revealed that single base spacer between 5mC-G steps is required for recognition of fully-mCHG through a dual flip-out of symmetric 5mCs from partner strands (Fig. 2B-C). Major difference in recognition through dual-flip out mechanism by UHRF1 SRA compared to other SRAs is that the bases from adjacent base pair are flipped-out in SUVH5 SRA-fully-mCG (5mCs; PDB ID: 3Q0C), SUVH5 SRA-hemi-mCG (5mC and C; PDB ID: 3Q0D) and UHRF2 SRA-hemi-hmCG (5hmC and C; PDB ID: 4PW6) complexes (Rajakumara et al. 2011; Zhou et al. 2014) (Fig. 3 A-C), and complementary bases are flipped-out in SUVH5 SRA-mCHH (5mC and G; PDB ID: 3Q0F), whereas in UHRF1 SRA-fully-mCHG, a base spacer is present between the flipped out 5mCs of both strands. It is intriguing that though UHRF1 SRA shares significant sequence (74%) and structural identities (RMSD:0.75Å) with UHRF2 SRA, in the context of hemi-CG sequence (hemi-mCG or hemi-hmCG), former adapts single base (5mC) flip-out mechanism, in contrast, dual bases (5hmC and C) are flipped-out in later case (Arita et al. 2008; Avvakumov et al. 2008; Hashimoto et al. 2008; Zhou et al. 2014).

**Fig. 3:**
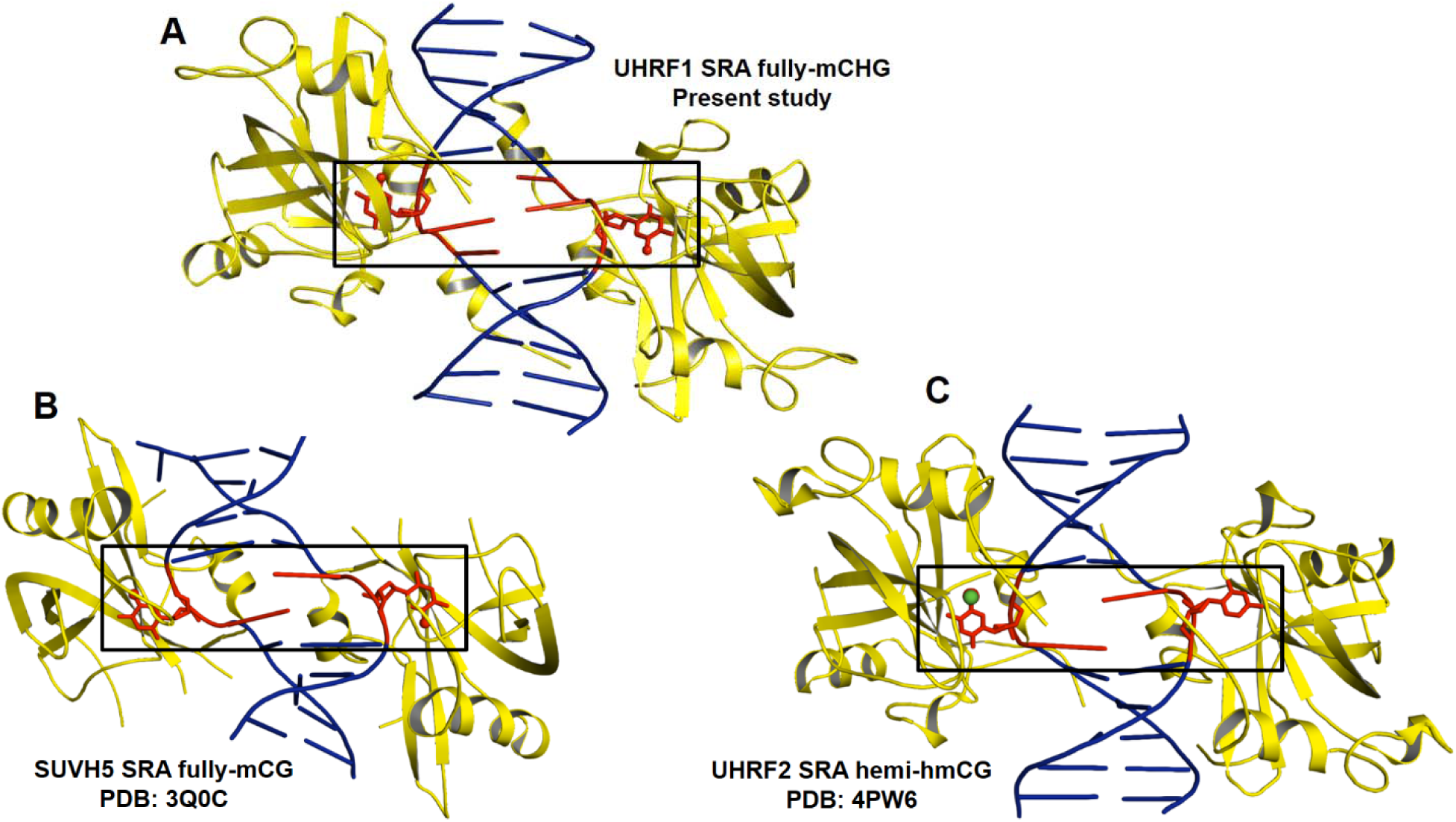
Structural comparison of SRA domains dual flipping out 5mC and other bases in different sequence context. **(A)** Flip-out of symmetrical 5mCs in UHRF1 SRA-fully-mCHG complex structure **(B)** Flip-out of symmetrical 5mCs in SUVH5 SRA-fully-mCG complex structure **(C)** Flip-out of 5hmC and Cytosine in UHRF2 SRA-hemi-hmCG complex structure. Base flipped out region of duplex DNA is highlighted in rectangle box. Protein (yellow) - DNA (blue) complex is represented in cartoon format except the flipped out base represented in red sticks.

### NKR finger is unstructured to recognize fully-mCHG complex by UHRF1 SRA

We analyzed the structure of NKR loop across the SRA domain of different proteins complexed with DNA containing a modified cytosine base to understand its role in 5mC/5hmC recognition in the context of different sequence (CG and CHG) and modification statuses (hemi or fully). NKR loop has been found to be unstructured in following complexes: UHRF2 SRA complex with hemi-hmCG (Zhou et al. 2014); SUVH5 SRA complex with fully-mCG, hemi-mCG, mCHH and fully-hmCG (Rajakumara et al. 2011, 2016) and SUVH6 SRA complex with hemi-mCHG (Li et al. 2018). Also, the NKR finger was found to be unstructured in UHRF1 SRA domain without DNA (Arita et al. 2008; Avvakumov et al. 2008) (Fig. 4). In contrast, it’s not only structured but also play a critical role in recognition of hemi-mCG DNA by UHRF1 SRA through interacting with orphaned ‘G’ in the duplex DNA and acts as selectivity filter by preventing the symmetry-related ‘C’ base from flipping out of the DNA duplex (Arita et al. 2008; Avvakumov et al. 2008; Hashimoto et al. 2008). Therefore, these comparative structural analyses indicate unequivocal role of NKR finger in recognition of hemi-mCG by UHRF1 SRA. NKR finger in UHRF1 SRA-fully-mCHG DNA (present study) is also unstructured suggesting that this region is highly flexible or dynamic in nature. The dynamic behavior of NKR finger may allow it to adapt different conformations to recognize cytosine methylation or hydroxy-methylation through dual-flip out mechanism (Rajakumara et al. 2011, 2016; Zhou et al. 2014).Otherwise, NKR fingers of the two SRA molecules involved in dual-flip out recognition may have steric clashes (Rajakumara et al. 2011, 2016; Zhou et al. 2014) (Supplementary Fig. 1). We speculate that flexibility of NKR finger would play a critical role in scanning both strands of the duplex DNA simultaneously for dual flip-out recognition of ‘cytosine’ methylation or hydroxyl-methylation in varied sequence context; but has a significant role in recognition of hemi-mCG byUHRF1 SRA (Zhou et al. 2014; Arita et al. 2008; Hashimoto et al. 2008; Avvakumov et al. 2008).

**Fig. 4:**
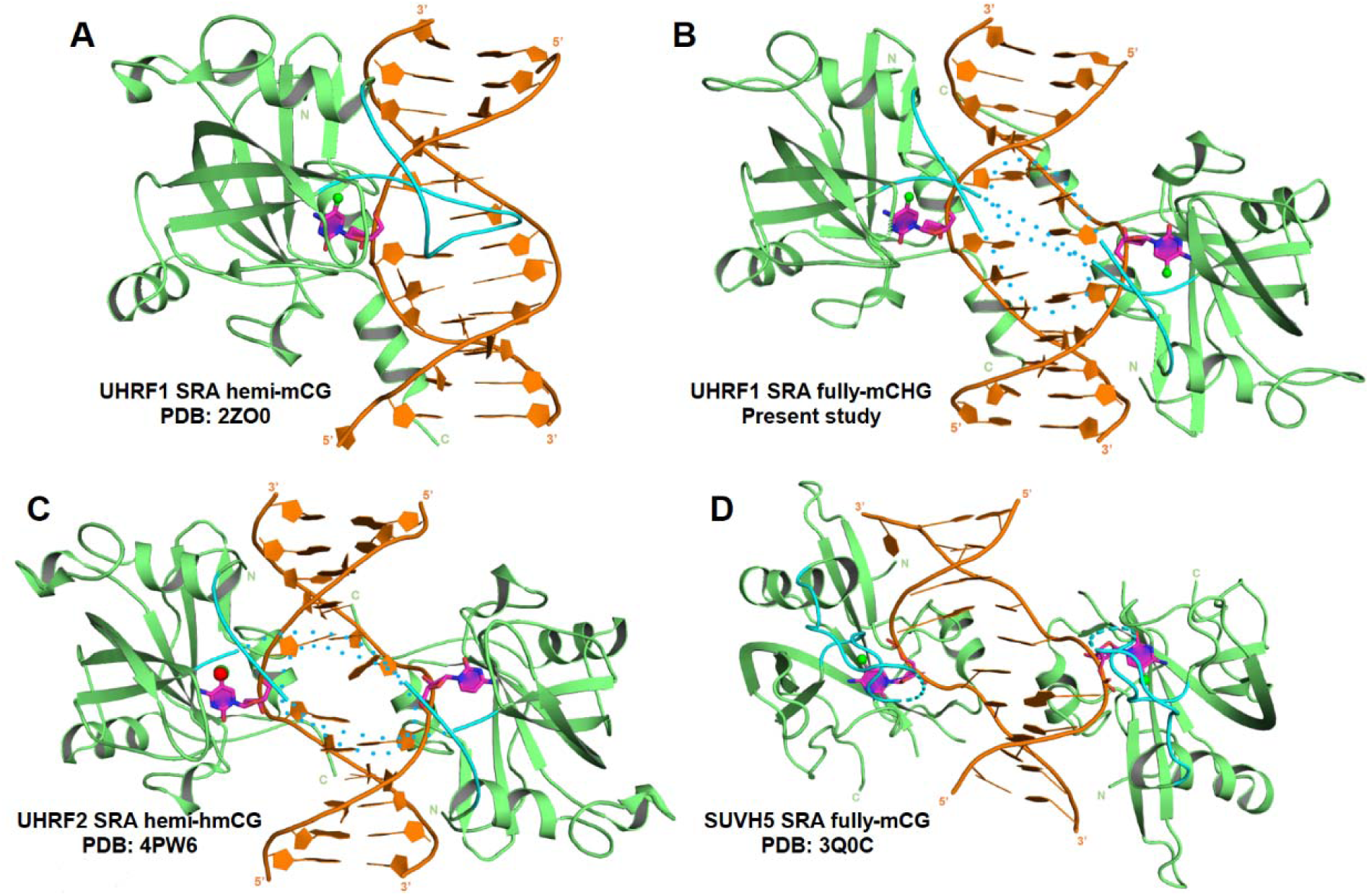
Structural comparison of finger loop among SRA domains. **(A)** UHRF1 SRA-hemi-mCG. **(B)** UHRF1 SRA-fully-mCHG **(C)** UHRF2 SRA hemi-hmCG. **(D)** SUVH5 SRA fully-mCG. Finger loop in all the structures is colored in cyan. Structured and unstructured finger loop is represented as solid and dotted loops, respectively. Protein (green) - DNA (brown) complex is represented in cartoon format. Flipped out base is colored in magenta. All the DNA bases are represented as rings.

## Conclusion

Our findings demonstrate an unanticipated versatility within UHRF1 SRA that recognizes 5mC not only in CG DNA context but also in non-CG DNA (mCH and fully-mCHG) context. Structural comparison among UHRF1 SRA-methylated DNA complexes revealed strikingly different mechanisms to recognize methylated DNAs: single flip-out for hemi-mCG and dual flip-out for fully-mCHG. Based on binding affinity and stoichiometry analyses we speculate that UHRF1 SRA may not form stable complex with fully-mCG DNA. Comparative binding and structural studies indicate that NKR finger has specific role in recognition of hemi-mCG through single flip-out mechanism and may contribute negligibly for binding affinity. Given the versatility of UHRF1 SRA for recognition and maintenance of different non-CG methylated DNA, and presence of non-CG methylation in brain neurons and germline cells, we infer that UHRF1 may have biological role in epigenetic regulation of non-CG DNA methylation and hence cell specific functions, which needs to be explored further.

## Methods

### Protein expression and Purification

The cDNA encoding full-length mouse UHRF1 was used for sub-cloning of SRA domain (amino acid residues 417-628) in pET28a-SUMO vector to generate a hexahistidine-SUMO tagged construct. UHRF1 SRA was expressed in *E*. *coli* Rosetta2 (DE3) (Novagen). For primary culture, freshly transformed colony was inoculated in LB broth and incubated overnight at 37°C. Primary culture (1%, v/v) was inoculated into secondary cultures and incubated at 37°C. After the OD_600nm_ reached 0.5-0.6, temperature was decreased to 18°C and the cultures were induced with 0.2mM of isopropyl-1-thio-Dgalactopyranoside (IPTG). At this temperature, the culture was incubated for 18-20 hours, following which the cells were harvested and re-suspended in lysis buffer (50mM Tris-HCl, pH 7.5, 150mM NaCl, 10mM imidazole and 3mM β-mercaptoethanol). Cells were lysed by ultrasonic homogenizer and then the lysate was clarified by centrifugation at 40,000g for 1 hour. The hexa-histidine-SUMO-tagged fusion protein was purified using a nickel charged column (HisTrap HP, GE healthcare). After elution with a 750mM imidazole containing buffer, the fusion protein was treatedwith Ulp1 protease at 25 Uml^-1^ to remove hexahistidine-SUMO tag, during a 16-hour dialysis using dialysis tube (10kDa cutoff), at 4°C. The protein was further purified by gel filtration chromatography (HiLoadSuperdex 75pg 16/600) pre-equilibrated with 25mM Tris-HCl, pH 7.5, 150mM NaCl, 3mMβ-mercaptoethanol). Purified UHRF1 SRA was concentrated to 25mgmL^-1^ at 4°C in 15mL, 3kDa cut-off Amicon(Merck-Millipore).

### DNA preparation

HPLC purified single stranded DNA oligos with 5mC and unmodified cytosine, along with complementary strands were purchased from W.M. Keck Foundation, Yale School of Medicine, Boston. The lyophilized single stranded oligos were dissolved in buffer comprising 10mM Tris (pH 7.5), 50mM NaCl and 3mM MgCl_2_. Concentration of each of the dissolved oligos were estimated and, self-complementary strand or equal concentrations of complimentary strands were mixed for annealing to form duplex DNA. The complex mixture was heated at 95°C for 5min and cooled at 4°C for overnight.

### Isothermal Titration Calorimetry (ITC) measurements

The molar equilibrium dissociation constant (K_D_), molar ratio (N) and thermodynamic parameters of the UHRF1 SRA bound to symmetric and asymmetric methylated non-CG DNA were determined using LV Affinity ITC (TA instruments) at 25°C. The buffer used for final gel filtration chromatography of the protein was used for diluting the DNA stock in order to minimize the difference in buffer composition of DNA and protein. The reference cell was filled with deionized water. The reaction cell was filled with 350μL of DNA (10-20μM) and sequentially titrated with 2μL injection of UHRF1 SRA (200-400μM) for a total 20 injections at 2 min interval. The data was processed using Nano-Analyze software. The ITC data was deconvoluted in “independent” model of curve fitting using a nonlinear least-squares algorithm. Binding enthalpy change (ΔH), association constant (K_A_), and binding stoichiometry (N) were permitted to vary during the least-squares minimization process. The errors denote standard deviations at 95% confidence interval.

### Crystallization

UHRF1 SRA-fully-mCHG complex was prepared by mixing purifiedUHRF1 SRA at a concentration of 3-10 mg/mL with fully-mCHG DNA duplex in a protein to DNA molar ratio of 1:0.6. The complex was incubated at 4°C for 30 minutes before setting up PEGRx crystallization screen (Hampton Research) using Mosquito robot (TTP Labtech). Crystallization screening was setupby mixing 300nL of protein-DNA complex solution with 300nL of reservoir solution in a 2 sub-well sitting drop (MRC-SD2) plate. The diffraction quality crystals of UHRF1 SRA-fully-mCHG DNA were grown in 0.1M Bis-Tris (pH 8.0), 0.2M Sodium Chloride and 12% PEG 3,350 at 18□C. The crystals were transferred to a cryoprotectant solution (reservoir composition with added 23% ethylene glycol) before mounting on to the goniometer head.

### Crystal data collection, structure determination and refinement

X-ray diffraction data from the UHRF1 SRA-fully-mCHG DNA complex crystals were collected at in-house X-ray source using Rigaku rotating anode generator and MAR345 image plate at CSIR-Centre for Cellular and Molecular Biology, Hyderabad, India. The UHRF1 SRA-fully-mCHG complex crystals diffracted to a maximum resolution of 3.0Å. The UHRF1 SRA-fully-mCHG complex crystals belonged to space group P2_1_2_1_2_1_. The images were processed using XDS and scaled using SCALA from the CCP4 program suite. The SRA domain structure of UHRF1 SRA hemi-mCG complex structure (PDB ID: 2ZO0) was used for phase calculation by molecular replacement method. Molecular replacement was done using MolRep in CCP4i suite. Electron density for all the bases of the DNA was unambiguous in the map generated using the SRA coordinates from Molecular replacement solution. The structure was refined using phenix.refine (Phenix suite). Model building was carried out using COOT (Emsley and Cowtan 2004). All the figures were prepared using PyMOL (DeLano, 2002). Refinement and model building were continued till R-work and R-free values converged. Data collection, refinement and Ramachandran statistics for the structure is given in Table 2.

## Acknowledgements

E.R. acknowledges Dr. Rajan Sankaranarayanan, Group Leader, CSIR-Centre for Cellular and Molecular Biology (CCMB). We thank Ms. Rukmini and Mr. Mallesham from CCMB for assistance in X-ray data collection. E.R. thanks the Department of Biotechnology (DBT) and the Science and Engineering Research Board, Department of Science and Technology, Government of India for the Ramalingaswami re-entry fellowship and Early Career Research Award, respectively. E.R. research is supported by extramural grant from the DBT and internal fund from the Indian Institute of Technology Hyderabad. N.K.N. and W.D thank University Grants Commission, Government of India for the fellowship. S.A. thanks the Ministry of Human Resource Development, Government of India for the fellowship.

