## Supplementary Fig. 1 for "UHRF1-SRA recognizes symmetric non-CG methylated DNA through dual-flip out of 5-methyl cytosines"

**Title**

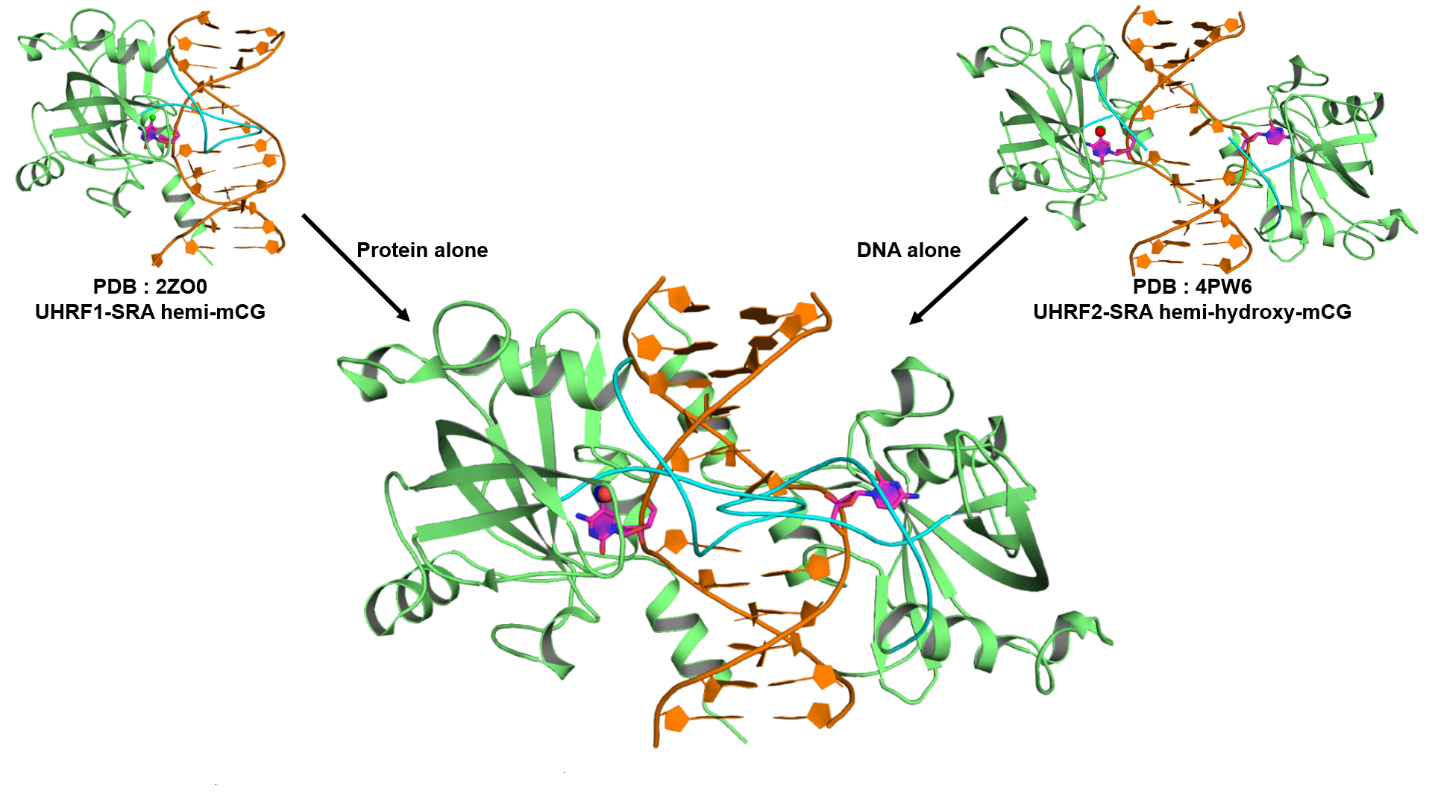

**Supplementary Fig. 1: Modelled structure showing the steric clashes of NKR loops (NKR loop is colored in cyan) from two SRA domains of UHRF1.** SRA domain from UHRF1 SRA-hemi-mCG complex (PDB ID: 2ZO0) is superimposed on the both the SRA domains of UHRF2 SRA-hemi-5hmCG complex structure (PDB ID: 4PW6) and the UHRF2 SRA domains are removed.The modelled complex consists of UHRF1 SRA recognizing hemi-5hmCG DNA with a dual flip out of 5-hydroxymethyl cytosine and cytosine bases (flipped out bases are colored magenta).
